# Corticocortical and Corticomuscular Connectivity Dynamics in Standing Posture: Electroencephalography Study

**DOI:** 10.1101/2024.04.30.591972

**Authors:** Kimiya Fujio, Kenta Takeda, Hiroki Obata, Noritaka Kawashima

## Abstract

Cortical involvements, including those in the sensorimotor, frontal, and occipitoparietal regions, are important mechanisms of neural control in human standing. Previous research has shown that cortical activity and corticospinal excitability vary flexibly in response to postural demand. However, it is unclear how corticocortical and corticomuscular connectivity is dynamically modulated during standing balance and over time. This study investigated the dynamics of this connectivity using electroencephalography (EEG) and electromyography (EMG). The EEG and EMG were measured in different 4 positions: sitting (ST), normal quiet standing (QS), one-leg standing (ON), and standing on a piece of wood (WD). For corticomuscular connectivity, we concentrated on sway-varying connectivity in the timing of peak velocity of postural sway in the anteroposterior direction. For the corticocortical connectivity, the time-varying connectivity was quantified, particularly in the θ-band connectivity which is linked to error identification, using a sliding-window approach. The study found that corticomuscular connectivity from the brain to the lower-limb muscle was strengthened during the sway peak in the γ- and β-frequency bands, while the connectivity strength from the muscle to the brain was decreased in the θ- and α-band. For the time-varying connectivity, the θ-connectivity in all time-epoch was divided into 7 states including both posture-relevant and -irrelevant clusters. In one of the 7 states, the strong connectivity pairs were concentrated in the mid-central region and the proportion of epochs from the ON and WD conditions was significantly higher, indicating a functional role for posture balance. These findings shed light on electrodynamic connectivity which varies in response to postural demand. Those dynamics, particularly in the θ-band connectivity, can be used for ongoing monitoring and/or intervention for postural disability.

## Introduction

Human standing is attributed to both the spinal and brainstem reflexes, as well as the cortical drives. Numerous studies have shown that these neural activities vary with postural demands for standing balance (Ouchi et al. 1999; Solopova et al. 2003; Fujio et al. 2018; Solis-Escalante et al. 2019). Transcranial magnetic stimulation studies indicate that the corticospinal excitabilities of the lower-limb muscles increase when standing balance is threatened (higher postural demand) by an unstable platform or mechanical perturbation (Solopova et al. 2003; Fujio et al. 2018). In addition, the pre-frontal and visual association areas activate more in challenging balance tasks, such as tandem posture or standing with eyes closed, using the hemodynamics approach (Ouchi *et al*. 1999; Mihara et al. 2008). The higher contribution of the sensorimotor cortex is thought to be beneficial for rapid recovery from a disturbance and maintaining upright posture.

Cortical oscillation is an effective probe for capturing posture-related brain activity. Electroencephalography (EEG) studies have shown that the power spectrum in the specific frequency, δ-, θ-, α-, β-, and γ-bands, can indicate the engagement of cortical drive (Thibault et al. 2014; Hulsdunker et al. 2015; Mierau et al. 2017; Fujio et al. 2023; Stokkermans et al. 2023). Higher β- and γ-powers appeared over occipital and frontal areas in upright posture compared to lying (Thibault *et al*. 2014), while α-power decreased at electrodes over the sensorimotor area before predictable perturbation (Solis-Escalante *et al*. 2019; Fujio *et al*. 2023). In the θ-frequency band, higher θ-power in the frontocentral area indicates error monitoring following the postural perturbation (Hulsdunker *et al*. 2015; Stokkermans *et al*. 2023). The change in each spectral power represents processing for a specific piece of information in response to postural demands.

Standing balance affects not only spectral power but also cortical and corticomuscular connectivity. Corticocortical θ- and α-networks are renewed in the frontoparietal and occipitoparietal regions when changing from a stable surface to an unstable surface for single-leg standing (Mierau *et al*. 2017), potentially leading to the facilitation of somatosensory processing and motor execution. A study using the postural perturbation paradigm found that a visual perturbation weakened α-cortical connectivity in the occipitoparietal area due to compared to the baseline activities before perturbation onset, while mechanical perturbation strengthened θ-connectivity in the central area (Peterson and Ferris 2019). Furthermore, the causal analysis found that mechanical perturbation increased corticomuscular θ-connectivity between various cortical areas, including anterior cingulate, parietal, and occipital areas and lower leg muscles. The θ-band connectivity improves in both ascending and descending directions, while the α-band connectivity decreases from the sensorimotor area to the lower muscles. Thus, corticocortical and -muscular connectivity are modulated both contentiously and transiently in response to postural demands for stable balance.

Although previous research has shown that corticocortical and corticomuscular connectivity strength varies with challenging postural tasks, little is known about its dynamics during normal standing. We suspect that this connectivity is modulated by spontaneous postural sway and time transition. The sway-varying connectivity is a posture-related change following the timing of the sway initiation and would be strengthened similarly to the response to an external perturbation. The time-varying connectivity is a sequential change over time and independently of postural events, as seen in resting-state brain dynamics (Allen et al. 2018; Wirsich et al. 2020). Given that the brain states fluctuate even at rest in a lying position (Deligianni et al. 2014; Allen *et al*. 2018; Wirsich *et al*. 2020), we hypothesized that cortical connectivity modulates over time in maintaining standing balance. Unraveling these questions will allow us to better understand the neural mechanisms that govern human standing and develop new treatments for postural disorders.

The purpose of this study was to clarify the dynamics of corticocortical and corticomuscular connectivity about standing balance. We concentrated on the impact of spontaneous posture sway and time transition on connectivity dynamics. Based on the above previous research (Mierau *et al*. 2017; Peterson and Ferris 2019; Fujio *et al*. 2023; Stokkermans *et al*. 2023), we inferred that both connectivity would be facilitated by the increased postural demand. Corticomuscular connectivity strength was measured at a specific phase in the anteroposterior (AP) sway in which the lower-limb muscles respond (Masani et al. 2003). Because of the strong relationship between postural sway and electromyography (EMG) activity, causality analysis was used to determine the information transfer separately for descending and ascending communication. It is critical to capture the appropriate bands for motor command and sensory input, respectively (Riddle and Baker 2005), and we considered that this connectivity relies on the γ- and β-frequency bands, where execution of voluntary movements is processed (Kilavik et al. 2013; Ulloa 2021). For corticocortical connectivity, we investigated the time-varying connectivity using a sliding window approach. Each epoch was examined, regardless of postural events using this approach. We considered that the increased cortical connectivity on the front-central-parietal region would emerge in more challenging tasks, particularly in the θ- and/or α-frequency bands which contribute to error monitoring and sensory integration (Sipp et al. 2013; Mierau *et al*. 2017; Stokkermans *et al*. 2023).

## Methods

### Participants

Seventeen adults participated (age: 29.2 ± 1.3 years old, height: 167.1 ± 2.1 cm, weight: 62.1±3.3 kg, 3 females). Exclusion criteria included a history of neurological disease, motor deficit, and current medication use. Participants were given a detailed explanation of the study protocol and gave their informed consent before the experiment. The experimental procedure was authorized by the ethical committee of the Research Institute of the National Rehabilitation Center for Persons with Disabilities. All experiments followed the principles outlined in the Declaration of Helsinki.

### Experimental procedures

The EEG, electromyography (EMG), motion capture, and grand reaction force data were collected in 4 different postures: sitting on a stool (ST), quiet-normal standing (QS), one-leg standing (ON), and standing on a piece of wood (WD, 7.5×42.0×7.5 cm). In the QS condition, the foot positions were predetermined to be 10 cm apart on both sides, parallel on a force plate. In the WD condition, a piece of wood was placed on the force plate and the foot positions were set as same as that in the QS condition, except that the support surface was narrower in the AP direction. In the ON condition, the right-side lower limb was predetermined as the supporting leg in all measurements, while the left-side hip and knee joints were held in a flexed position without contact with the supporting leg. Participants were instructed to maintain their gaze on a visual target 1.5 m in front of them without making any deliberate movements. Before the experiment, a practice session was held to ensure that the necessary postures were maintained. The order of the experimental conditions was chosen at random, and each condition received 8 trials. Each trial was timed for 60 seconds.

### Data acquisition

The EEG signals were collected at 1,000 Hz with a 32-electrode system (g.LADYbird active, g.tec, Austria). The electrode placement was based on the international 10–10 system. The reference and ground electrodes were attached to the right and left mastoid processes, respectively. Electromyographic (EMG) activities were measured from 6 muscles, including the bilateral tibialis anterior (TA), soleus (SOL), and medial gastrocnemius (GA) muscles, using a wireless EMG system (Trigno, Delysis Inc., USA, sampling rate: 1,000 Hz, bandpass filter: 15-750 Hz). The EMG sensors were securely attached to the muscle belly and wrapped in thin elastic bandages. Body kinematics were recorded using a motion capture system equipped with 13 infrared cameras (Mac3D system, Motion Analysis, Corp, USA, sampling rate: 200Hz). We used a Helen Hayes marker set consisting of 29 reflective markers attached to body landmarks. The positions’ time series were filtered with a fourth-order, zero-phase-lag Butterworth low-pass filter with a cutoff frequency of 6 Hz.

Grand reaction forces were collected at 1,000 Hz from a single force plate system to obtain the center-of-pressure (COP) coordinates (9281B11, Kistler Instrument AG., Switzerland). The force data was filtered using a fourth-order, zero-phase-lag Butterworth low-pass filter with a cutoff frequency of 10 Hz. The EEG, EMG, motion, and force data were aligned using a common signal and treated as a single multivariate dataset during the subsequent preprocessing.

### Data analysis

#### EEG preprocessing for single-subject

The EEG data was preprocessed using EEGLAB (Delorme and Makeig 2004), which is included as a toolbox for MATLAB (Mathworks Inc., USA). Figures 1A and B depict the outline of EEG processing. The EEG data were high-pass filtered, with a zero-phase 1.0 Hz cutoff, and then down-sampled to 200 Hz. After reducing line noise at 50 Hz with the cleanLineNoise function in the EEGLAB plugin, we removed bad channels with obvious artifacts through visual inspection. Artifact subspace reconstruction was used on the remaining data to remove flat channels and high-amplitude artifacts (time length for flat channels: 5s, channel correlation for threshold: 0.85, standard deviation for threshold: 10, sliding window: 500 ms). The continuous EEG data was segmented into 5s epochs for automatic rejections (exceeding the average±5 standard deviation) and re-referenced to the common average. An adaptive mixture independent component analysis was applied to the concatenated EEG to distinguish between brain and non-brain components and to identify the cortical source. Finally, a single equivalent current dipole was calculated for the scalp projection of each independent component (IC) using a standard MNI model.

**Figure 1.**
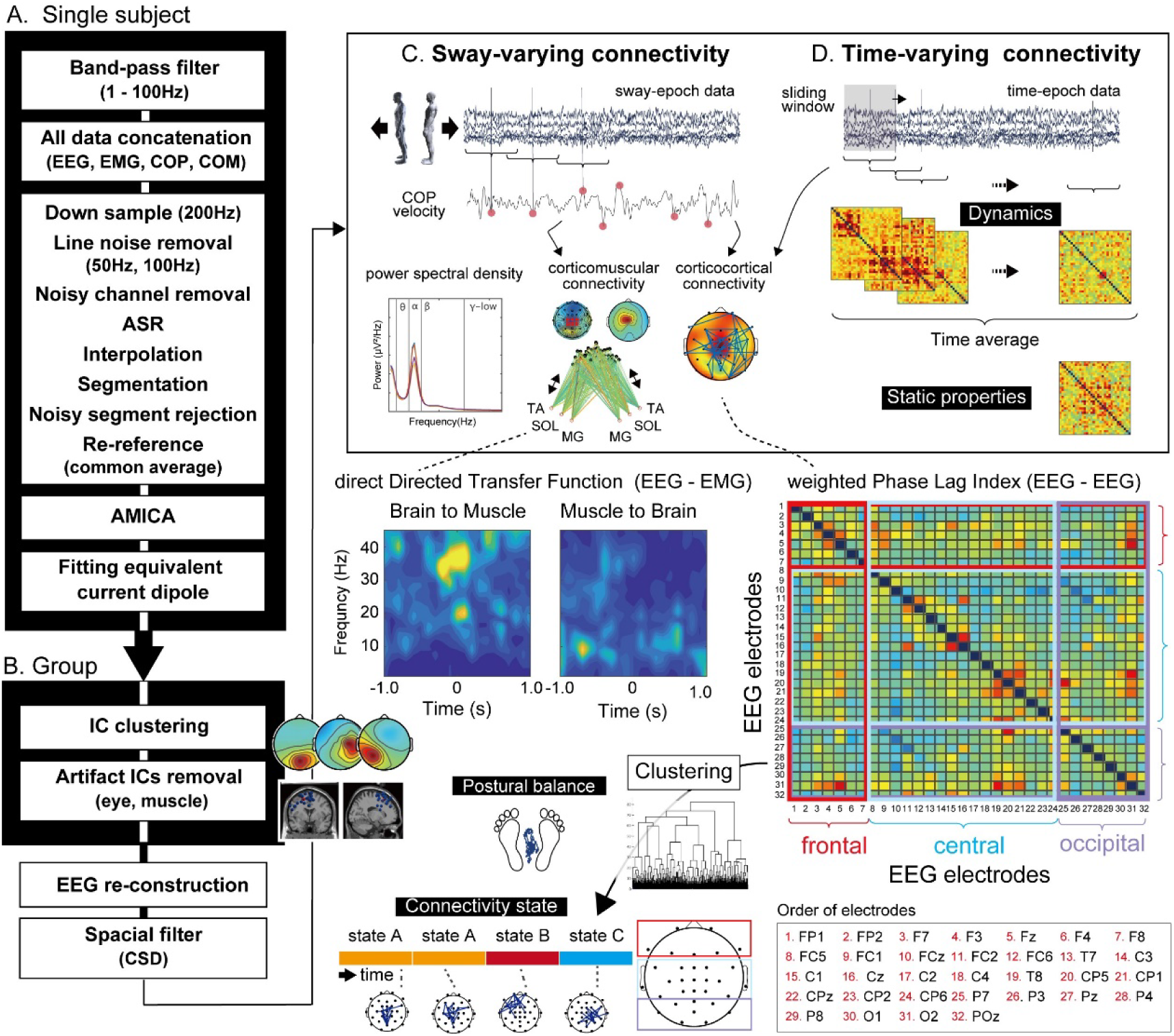
Overview of EEG-processing pipeline. The left column depicts the analysis flow for single-subject and group data (A and B). The cleaned EEG was segmented into sway- and time-epoch (C and D) before being used to calculate the corticomuscular and/or corticocortical connectivity and power spectral density. The Dynamics focused on in this study are sway-varying corticomuscular and time-varying corticocortical connectivity. Static properties mean the averaged feature of cortical connectivity in all time-epoch data. The corticomuscular connectivity strength (dDTF) was quantified using a causality analysis. For time-varying cortical connectivity, all epochs from all conditions were classified into the subcategory for connectivity state using the hierarchical cluster analysis with feature vectors from all pairwise wPLI. Standing balance was finally compared among different clusters.

#### IC clustering for all-subject group

To categorize ICs from all participants based on their similarities, clustering analysis with robust k-means ++ was used. The ICs for clustering were retained when the residual variance was less than 15%. The robust k-means ++ iterated 500 times with a three-dimensional dipole location as the feature vector, omitting the outliers that were above 5 SD from the other cluster centers. The number of clusters was determined to be 13 after evaluation using the gap statistics. According to previous research (Wagner et al. 2016; Artoni et al. 2017), the artifact IC cluster associated with eye blinks or movements was identified based on the topographic power map, distribution of the power spectrum, and dipole location. We further classified the scalp projection of each IC using the ICLabel function implemented in the EEGLAB plugin and removed non-cortical ICs that were identified as “Muscle” and “Eye” components based on their power spectrum, topographic power map, and dipole position. Finally, the EEG signals were reconstructed by back-projecting cortical ICs and applied to a spatial filter, current source density transformation (Perrin et al. 1989; Kayser and Tenke 2006), to reduce the volume conduction effect (order of spline = 4, maximum order of the Legendre polynomial = 50, precision = 10^−5^).

#### Head kinematics and postural sway

To identify excessive head movements that may cause EEG artifacts, the trajectory of a center-of-mass of the head segment (COM_head_) was calculated from a time series of 3 marker positions (frontal, parietal, and rear head). We further differentiated it to determine the velocity of COM_head_. In addition, the COP velocity (COPv) was calculated by differentiating the time series of COP coordination to detect peaks of postural sway in the anteroposterior direction and assess postural stability.

#### Data epochs

In this study, we focused on posture sway- and time-varying neural states, as well as their static properties while standing quietly (Figure 1C and D). To this end, the time series of concatenated data was divided into epochs using two methods: the sway-detection and the time-sliding methods. The sway-detection method identified negative and positive peaks of the COPv in the AP direction. It was previously known that compensatory EMG activity is elicited in the ankle muscle with little delay following the COPv peak (Masani *et al*. 2003). To avoid searching for consecutive points in a short time, we set a 2.0-second interval in which no peak points were searched after a previous peak. Based on these peak timings, time series data were extracted with a window of 1.5 seconds before and after the peak timing (sway-epoch data, Figure 1C). In the time-sliding method, we separated the entire length of data using a rectangular window that was 8.0 seconds long and overlapped by half that length (time-epoch data, Figure 1D). To reduce the impact of excessive head movements on EEG data, we rejected epochs where the velocity of COM_head_ exceeded the averaged root-mean-square ± their two-standard deviation in sway- and time-epoch data, respectively.

To obtain static properties of corticocortical connectivity in each posture, we averaged over all epochs extracted using the time-sliding method. It indicates a common trend of the modulation of connectivity strength due to posture changes, which can confirm the findings in the previous research (Hulsdunker *et al*. 2015; Mierau *et al*. 2017; Varghese et al. 2019).

#### Corticomuscular connectivity

Using the sway-epoch data, we investigated the causal interaction for corticomuscular connectivity between the sensorimotor region and lower-limb muscles. To estimate the direction of information transfer, we used spectral analysis with the multivariate autoregressive (MVAR) model. One of the 6 EEG electrodes, Cz, C1, C2, CPz, CP1, CP2, and bilateral TA, SOL, and MG, were combined into a single dataset. The EMG signal was high-pass filtered at 2 Hz (3rd-order Butterworth filter), detrended, and normalized for time average and standard deviation. For the MVAR model fitting, a time window was taken for 400 ms with a step size of 100 ms. The model order was set to a value of 15 (75ms for 200 Hz of the sampling rate), which is slightly larger than the model for upper motor tasks (Brovelli et al. 2004) due to the longer lag between the sensorimotor cortex and the lower limb muscles. We also checked the optimal model older against the Akaike Information Criteria before fitting the MVAR model using the Vieira-Morf algorithm in each participant.

A connectivity strength was determined by the direct Directed Transfer Function (dDTF) which is a causal estimator in the frequency domain (Korzeniewska et al. 2003). The dDTF is calculated as the product of the full-frequency DTF and partial coherence and thus is resistant to the effects of indirect transmission. We calculated the dDTF at 1–50 Hz for the descending (brain to muscle) and ascending (muscle to brain) directions, respectively. The analyzed window was further divided into 3-time ranges based on the compensatory EMG activation timing (−900–−300 ms, −300–300 ms, 300–900 ms). The dDTF were averaged in 6 pairs from each EEG electrode and compared across 4 frequency bands (θ, α, β, γ). In this study, we focused specifically on the dDTF to the compensatory activated muscles: the MG and SOL in the anterior sway and the TA in the posterior sway. To validate the results of this sensor space analysis, the same procedure was repeated using motor IC (Supplementary Figure S1). We used the source information flow toolbox throughout the process (Delorme et al. 2011).

#### Corticocortical connectivity

To reveal posture-relevant global connectivity, we evaluated phase coupling for pairwise electrodes using the weighted phase lag index (wPLI) (Vinck et al. 2011):

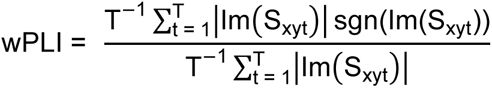

where *T* is the number of time points, *Im (S_xyt_)* represents the imaginary part of the cross-spectrum at time *t*, and *sgn* indicates the sign. The wPLI was calculated for each epoch of both sway- and time-epoch data for 4 frequency bands (θ-, α-, β-, γ-band), respectively. To investigate the effects of posture changes, the wPLI was converted into a robust z-score (wPLIz) in all conditions. Before performing the following principal component analysis (PCA) on the time-epoch data, the wPLI was normalized in all participants.

#### Clustering for the time-varying corticocortical connectivity

To estimate the transitions of neural states in sitting and standing, time-epoch data was classified using hierarchical cluster analysis. The connectivity at the θ-frequency band was used for the following analysis. First, a feature vector for classification was extracted from the wPLIz matrix of all participants using the PCA. After performing singular value decomposition of the matrix, the principal comportments (PC) scores and factor loading were calculated. The number of PCs was determined to be greater than 10% of the variance for the hierarchical cluster analysis. The Euclidian distance was calculated between each pair, and Ward’s method, which determines the minimum variance of squared distance within a cluster, was used as a linkage criterion. The number of clusters was estimated using the upper tail method, and the dendrogram was drawn to confirm the discrimination of those clusters. Finally, the proportion of posture condition was calculated in each cluster to demonstrate the characteristics of each cluster, particularly their relevance to postural stability.

In addition, we considered that posture-specific connectivity states would be highlighted in the timing of sway peaks. Thus, we further examined whether the posture-relevant PC appeared more frequently in the sway-epoch data. To that end, the classifier was built using the time-epoch data and applied to the sway-epoch data. We used a support vector machine (SVM) with a non-linear radial basis function and the PC scores from the wPLIz matrix of the time-epoch data as the feature vector. The parameters C and Gamma were adjusted using grid search. For a performance test, 5-fold cross-validation was performed to calculate the confusion matrix. The accuracy, precision, recall, and F1 score were calculated to validate of the SVM model. Finally, the learned SVM model was used to classify the wPLIz in the sway-epochs and determined the frequency of each state in anterior and posterior sway, respectively.

#### Power spectrum density

To characterize the EEG spectral power for each electrode, the power spectral density (PSD) was calculated using Welch’s method at each epoch. After removing the linear trend, a fast Fourier transformation of the EEG waveform was performed with an overlap of 50% window length. A Hamming window was used to reduce edge effects. The PSD of all epochs was averaged in each participant for sway-varying connectivity and in all participants for time-varying connectivity.

### Statistical analysis

To assess cortico-muscular connectivity, dDTF was measured at 3 time points (T1: −900–−300 ms, T2: −300–300 ms, T3: 300–900 ms) and averaged across the 6 electrodes. A two-way ANOVA was used to confirm the effects of POSTURE and TIME on corticomuscular connectivity. When sphericity could not be assumed in Mendoza’s multisample sphericity test, the variables were adjusted using the Greenhouse-Geisser correction. When an interaction was significant, simple main effects were investigated at each level. In the post-hoc comparisons, Shaffer’s post-hoc tests were used to account for differences between 3 standing postures, except for the ST condition, and 3 time intervals. We further quantified the dDTF using the IC allocated in the sensorimotor cluster to demonstrate the validity of the sensor-level analysis.

For the corticocortical connectivity, we analyzed wPLIz between 496 electrode pairs using the one-way ANOVA to determine the effect of POSTURE. False discovery rate (FDR) correction was used for post-hoc comparison across all pairs. A one-way ANOVA was also used to compare the PSD of all electrodes between different postures or states, and the FDR correction was used for the post-hoc tests.

To compare the proportions of 4 postural conditions in each cluster, we used the one-way ANOVA with Shaffer’s test for the post-hoc comparisons. Furthermore, to compare the allocation of each state in the time- and sway-epoch data, the independence of distribution was tested using a chi-squared test.

To investigate the relationship between corticocortical connectivity and standing balance, integrated EMG activity (iEMG), background EMG activities (BGA), and COPv were compared across standing 3 conditions and different clusters. The iEMG was calculated from the rectified EMG waveforms between −300–300 m after the above preprocessing and subtracted from the integrated BGA. The BGA was defined from −500ms–−300ms which was the period just before EMG elicitation. The iEMG was finally normalized using the highest value among the 3 standing conditions in each participant. One-way ANOVA for the iEMG and BGA, as well as the Kruskal-Wallis test for the COPv, were used, and, if the main effect was significant, Shaffer’s multiple comparisons were applied. For all comparisons, the significant level was set at α = 0.05, and the q-value for FDR was set at 0.1.

## Results

Thirteen common IC clusters were identified from the EEG time series in 4 different postures (Supplementary Figure S1). A total of 50 individual ICs were removed as noise components. After removing the artifacts ICs, the connectivity analysis was performed using the cleaned EEG. For the sway-varying connectivity, the average number of epochs was 50.7 ± 0.8 in 3 conditions and both anterior and posterior directions (anterior, QS: 47.3 ± 1.7, ON: 52.9 ± 1.9, WD: 51.6 ± 2.2 epochs; posterior, QS: 51.3. ± 1.8, ON: 50.7 ± 1.9, WD: 50.4 ± 2.1 epochs). For the time-varying connectivity, the sliding-window method was used to extract a total of 6997 epochs from all participants in all conditions.

### Sway-varying corticomuscular connectivity

Figure 2A depicts a typical time series of EEG, EMG, and COPv extracted from the peak of COPv during the AP sway. The time-frequency maps of EEG (Cz) and EMG (MG) that were used for the connectivity analysis were also shown. Both anterior and posterior epochs were segmented as the reference to the COPv peak. Note that the peak of MG muscle activation coincides with the peak of COPv when swaying anteriorly, while it is temporarily suspended in the case of posterior sway. The MG activation counteracts the gravity torque in the ankle joints to compensate for anterior tilt (Masani *et al*. 2003). There were no apparent changes in the time series of Cz. Figure 2B depicts the time series of compensatory EMG activities within the −1,000 ms–1000 ms window. As with the same of MG, the TA activation was evoked particularly in ON and WD conditions, which coincided with the peak of COPv in the posterior direction. We investigated whether both iEMG and BGA were different from postural conditions. A one-way ANOVA revealed a significant main effect of postural condition in the iEMG of MG in the anterior sway (F_1.66, 26.57_ = 23.12, p < 0.001, generalized eta^2^=0.44) and that of TA in the posterior sway (F_1.48, 23.74_=33.60, p<0.001, generalized eta^2^ = 0.5685). A post-hoc test revealed that MG activities increased in 2 standing conditions, particularly in the ON and WD conditions, whereas the TA activity increased significantly only in the WD condition. A similar trend was observed in the BGA: There were main effects in both anterior (MG, F_1.55, 24.79_ = 76.80, p < 0.001, generalized eta^2^ = 0.70) and posterior directions (TA, F_1.59, 25.43_ = 70.55, p < 0.001, generalized eta^2^ = 0.68). In the post-hoc tests, significant increases in the MG were observed in the ON and WD conditions while swaying anteriorly (QS vs. ON, p < 0.001, ON vs. WD, p = 0.002, ON vs. WD, p < 0.001). In the posterior sway, there were clear increases in the ON condition as well as in the WD condition (QS vs. ON, p < 0.001, ON vs. WD, p = 0.003, ON vs. WD, p < 0.001).

**Figure 2.**
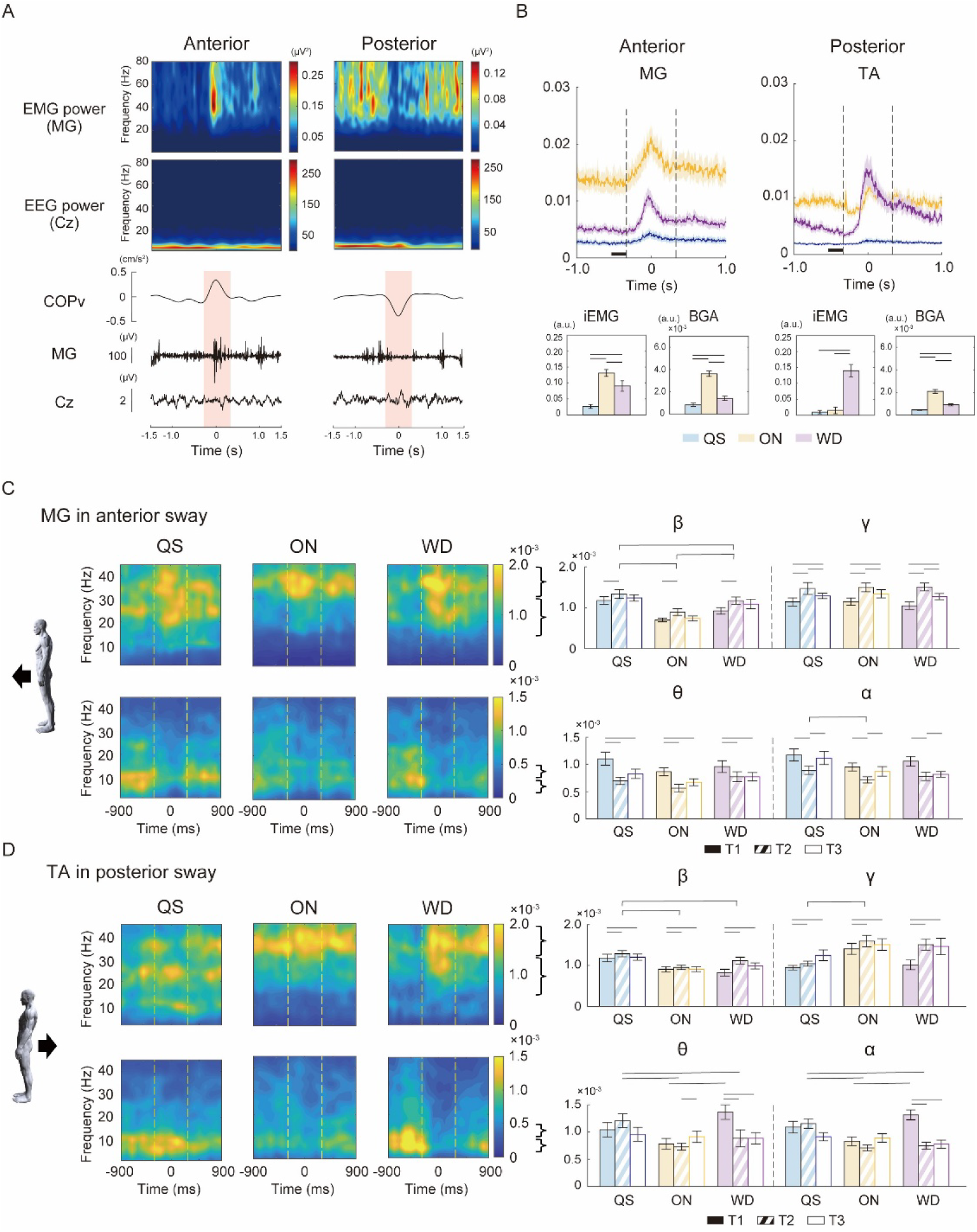
Tracking of the EMG activities and corticomuscular connectivity while postural sway. A. A typical time series of EEG (Cz), EMG in MG muscle and COPv based on AP direction of postural sway. The time-frequency EEG and EMG (MG) power patterns are shown. The transparent red window represents the analysis period: −300–300 ms. B. EMG activities during 3 standing conditions in the sway-determined epoch. The black square bar represents the timeframe for baseline activities. left; MG activity in anterior sway. right; TA activity in posterior sway. iEMG refers to the integrated EMG activity within the −300–300 ms window. BGA denotes background EMG activities. C. Time-frequency maps of corticomuscular connectivity (left) and mean strength in β- and γ-frequency bands for the descending communication and θ- α-frequency bands for the ascending communication, (right). For the anterior sway, the connectivity strength was depicted between the brain and the MG muscle (upper; the descending connectivity, lower; the ascending connectivity). For the posterior sway, the TA muscle was assessed for connectivity strength (D). The bracket represents a significant main effect on POSTURE. Dash bar means a significant difference in a post-hoc test.

Corticomuscular connectivity was modulated in different ways along ascending and descending pathways. Figure 2C and D display the time-frequency modulation of dDTF and their strength in each band and condition for the compensatory-activated muscle. Strong descending connectivity (EEG to EMG) was observed in the γ- and β-frequency bands to the θ- and α-frequency bands in both muscles. Thus, we particularly focused on the γ- and β-frequency bands for the effects of POSTURE and TIME in the descending connectivity. For the γ-connectivity in the MG (Figure 2C), a two-way ANOVA found a significant main effect on TIME (F_1,98 31.7_ = 15.01, p < 0.001, generalized eta^2^ = 0.713) and no interaction (p = 0.794). The post-hoc comparisons revealed increased connectivity at T2. The β-connectivity had significant effects on TIME (F_1.82, 29.18_ = 7.25, p = 0.003, generalized eta^2^ = 0.05) and POSTURE (F_1.6, 25.68_ = 22.60, p < 0.001, generalized eta^2^ = 0.24) without their interaction (p = 0.744). Post-hoc comparisons showed that β-connectivity was higher at T2 in each condition and larger in QS compared to ON and WD conditions. In posterior sway, the TA muscle showed similar facilitation (Figure 2D). Significant effects for the γ-connectivity were observed on TIME (F_1.76, 28.15_ = 7.28, p = 0.003, generalized eta^2^ = 0.05) and POSTURE (F_1.83, 29.27_ = 4.78, p = 0.018, generalized eta^2^ = 0.10) and no interaction (p = 0.067), with significant enhancements observed at T2 and T3, especially in the ON condition. The β-connectivity had significant main effects on TIME (F_1.85, 29.58_ = 4.90, p = 0.016, generalized eta^2^ = 0.041) and POSTURE (F_1.8, 29.9_ = 9.27, p < 0.001, generalized eta^2^ = 0.16) without their interaction (p = 0.165). Post-hoc comparisons showed a significant increase in the QS condition at T2.

For the ascending corticomuscular connectivity (EMG to EEG), strong connectivity was shown at the θ- and α-frequency bands (Figure 2C). A statistical test revealed a significant TIME (F_1.83, 29.29_ = 10.73, p = 0.004, generalized eta^2^ = 0.11) for the θ-band, and stronger connectivity was observed in the T1 than the other time-ranges in post-hoc comparisons. In the α-connectivity, the main effects were significant for both TIME (F_1.91, 30.53_ = 7.38, p = 0.027, generalized eta^2^ = 0.09) and POSTURE (F_1.45, 23.22_ = 4.69, p = 0.028, generalized eta^2^ = 0.06) with no interaction (p = 0.585). The post-hoc tests revealed stronger connectivity in QS but weaker strength at T2. In posterior sway, we found a significant interaction between θ-connectivity (F_2.89 46.20_ = 5.23, p = 0.003, generalized eta^2^ = 0.07) and simple effects (TIME: ON, F_1.74, 27.79_ = 4.02, p = 0.034, generalized eta^2^ = 0.04; WD, F_1.96, 31.33_ = 9.93, p < 0.001, generalized eta^2^ = 0.155; POSTURE: T1, F_1.92, 30.74_ = 7.42, p = 0.002, generalized eta^2^ = 0.19; T2, F_1.91, 30.57_ = 5.90, p = 0.007, generalized eta^2^ = 0.14). In the α-frequency band, there was a significant interaction (F_2.65 42.35_ = 8.49, p = 0.003, generalized eta^2^ = 0.14) and simple effects were revealed on both TIME (WD, F_1.37, 21.94_ = 23.67, p < 0.001, generalized eta^2^ = 0.41) and POSTURE (T1, F_1.79, 28.61_ = 5.58, p = 0.011, generalized eta^2^ = 0.21; T2, F_1.66, 26.56_ = 14.35, p < 0.001, generalized eta^2^ = 0.34, (Figure 2D).

The IC level analysis revealed a similar trend, validating the above sensor-level analysis (Supplementary Figure. S3). The distribution maps of normalized dDTF under all conditions were also shown to highlight the differences between conditions (Supplementary Figure. S4C).

### Time-varying corticocortical connectivity and power spectrum density

Figure 3 shows the results of clustering for time-epoch data. Based on the predetermined threshold of the variance, 7 PCs were extracted as a feature vector with a PCA of all 6997 epochs. Thus, a 7-dimensional feature vector was used for hierarchical cluster analysis. As a result, 6997 epochs were divided into 7 distinct clusters (State A-G, Figure 3A). Each cluster was distinguished by a major connection accounting for 5% of the top strength and the proportion of posture conditions. State A (11.2% in total) had a small PSD in the central area and a higher proportion of ST conditions. There is no significant connectivity in the central region. State B (7.5%) exhibits asymmetric connectivity, particularly in the right hemisphere and a small proportion of the ON condition. A large and widespread PSD was observed in the central area. State C (19.8%) has significant connectivity in the occipital area and a high prevalence of all conditions. State D (17.6%) is distinguished by its strong connectivity in the central area. Note that, in this state, the proportion was higher in ON and WD conditions compared to ST and QS conditions (F_1.78, 28.42_ = 9.25, p = 0.001, generalized eta^2^ = 0.23, Figure 3B). State E, F, and G (10.2%, 13.1%, 20.6%, respectively) have stronger connectivity in the frontal, left frontocentral, and right frontocentral areas. The proportion of these states was nearly equal across the 4 postures.

**Figure 3.**
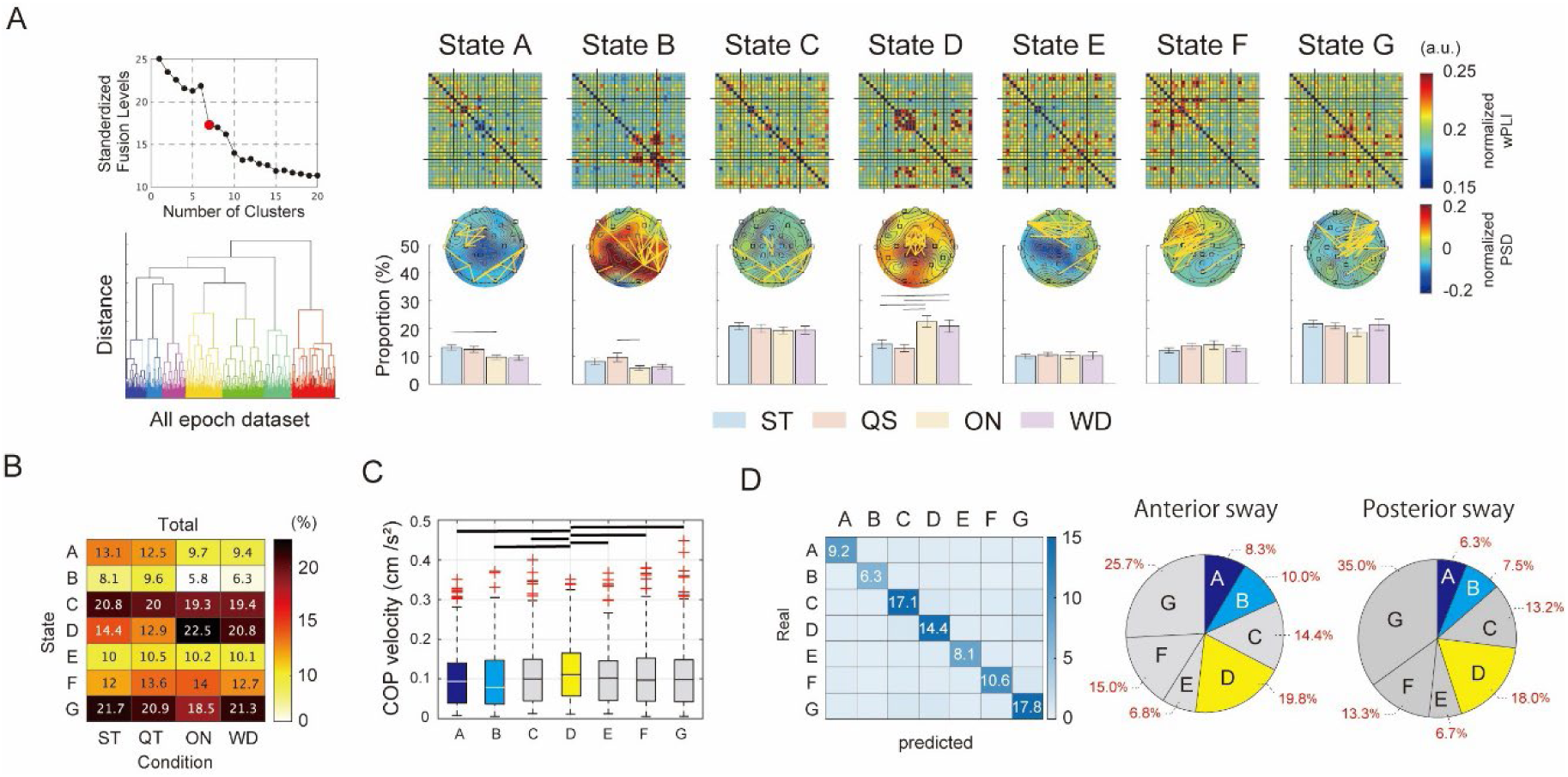
Time-varying corticocortical connectivity and its differences among postural conditions. A. Seven neural states were classified using hierarchical cluster analysis for time-varying corticocortical connectivity. The clustering order is determined by the dendrogram generated from the feature matrix. Left; standardized fusion levels for the number of clusters and the dendrogram. Right; connectivity maps, the top 5% strength of connectivity on topography, and the proportion of posture conditions in each state. B. Proportion of postural conditions in 7 states for all participants. C. Comparison of COP velocity among 7 states. D. The sway-epoch data is divided into 7 states using the SVM model. left; Confusion matrix for the SVM model based on 6997 time-epoch data. middle and right; Allocation of the sway-epoch data in the anterior sway (middle) and in the posterior sway (right).

### Functional relevance of the corticocortical connectivity for standing balance

To demonstrate the importance of cortical connectivity in standing balance, we compared the COPv across different states (Figure 3C). Statistical tests revealed a significant main effect on POSTURE (p<0.001, F_6, 52_=5.65, generalized eta^2^=0.006) and a higher COPv in State D compared to other clusters.

We also attempted to confirm the effect of postural stability using a classification approach with sway-epoch data. The accuracy of the SVM model using time-epoch data was 83.5% (precision: 83.2%, recall: 83.1%, F1 score: 0.83, Figure 3D). Using this model, we examined the distribution of the sway-epoch data across the 7 states. When the sway-epoch data was applied to this model, 19.8% was allocated to State D for anterior and 18.0% for posterior sway. The chi-square test revealed a significant result in all pairwise tests (time epoch vs. sway anterior, χ^2^ = 100.06, p < 0.001, time epoch vs. sway posterior, χ^2^ = 234.23, p < 0.001, sway anterior vs. sway posterior, χ^2^ = 50.93, p < 0.001, (Figure 3D). Thus, the occurrence rates are not independent of time- and sway-epoch data, indicating that State D is not the majority even in the timing of sway peak. The posture-related changes in COPv, PSD, and corticocortical connectivity in the sway-epoch data were shown in Supplementary Figure S5.

### Static properties of corticocortical connectivity and the power spectrum density

Figure 4A shows the significant pairs of corticocortical connectivity averaged across all windows in each condition. The statistical analysis revealed that corticocortical connectivity is reorganized based on postural states, particularly at the θ-, α-, and β-frequency bands. The high connectivity at the θ-band localizes mainly in the frontal and central areas, and their strength tends to be increased during ON and WD conditions. Contrary, α-connectivity strength increased more in ST and QS conditions compared to ON and WD conditions. In the β-frequency band, topography shows similar significant pairs to the α-band, but connectivity strength on the map differs in WD.

**Figure 4.**
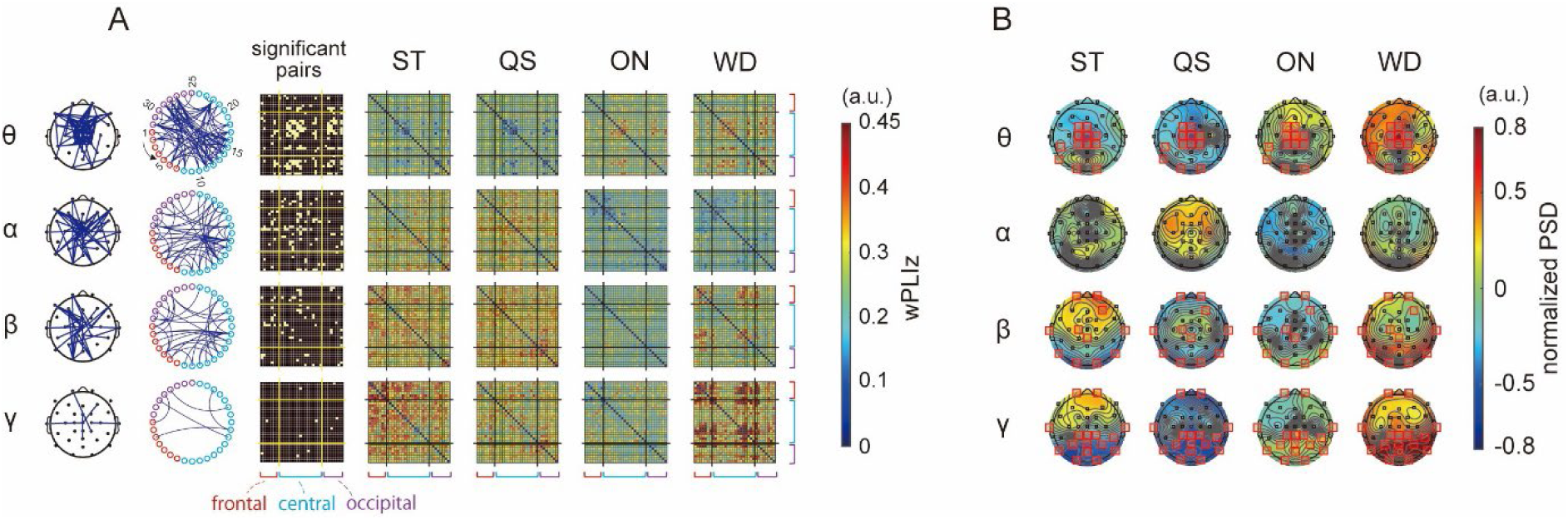
Static properties of corticocortical connectivity and PSD. A. Time-averaged wPLI for each frequency band and 4 conditions. The significant pairs across conditions are represented by topographies, circular graph diagrams, and matrices. Connectivity strength (wPLIz) is also shown for all pairs. The numbers on the circular graph correspond to the electrodes in Figure 1. B. Time-averaged PSD for each frequency band and 4 conditions. Red squares and black circles indicate the significant and non-significant electrodes, respectively.

Figure 4B depicts the distribution of averaged PSD and significant electrodes in one-way ANOVA across postural conditions on the topography. In the θ-frequency band, significant electrodes are concentrated around the central midline region, with higher power in the ON and WD conditions compared to ST and QS conditions. In γ-frequency bands, the significant differences are primarily observed in the occipital and parietal areas. In the β-frequency bands, electrodes are distributed more widely, including the central, occipital, and central regions. In addition, the static differences of the COPv showed a main effect (p < 0.001, F_1.30, 20.78_ = 77.02, generalized eta^2^ = 0.58) and significantly increased in the WD condition (QS vs. ON: p < 0.001, QS vs. WD: p < 0.001, ON vs. WD: p = 0.013).

## Discussion

This study aimed to show the modulation of corticocortical and corticomuscular connectivity due to time transition and postural sway. We hypothesized that postural demands would further affect both connectivity. The results showed a transient increase in corticomuscular connectivity between the sensorimotor region and the lower muscles while swaying in both anterior and posterior directions. The dominant frequencies vary depending on the transfer direction: β- and γ-frequency bands for the descending connectivity, θ- and α-frequency bands for the ascending connectivity. We found the strength β-connectivity in QS and WD and the weaken α-connectivity in ON and WD conditions, suggesting the influence of postural demands on the modulation. In addition, focused on the θ-frequency band, the time-varying cortico-connectivity for 4 different postures was classified into 7 states including posture-relevant components. These findings shed light on the dynamics of neural connectivity during postural control and support the evidence for the distinct functional role in the frequency bands.

### Sway-varying corticomuscular connectivity

We found that corticomuscular connectivity significantly changes depending on the phase of postural sway and posture conditions. The dominant frequency of this modulation differed between ascending and descending communications, with β- and γ-bands for descending and θ- and α-band for ascending connectivity. The descending connectivity increased following compensatory muscle activation, while the ascending connectivity decreased at the same time. Previous studies suggest that γ-oscillations play a role in motor execution of forearm and finger movements (Brown et al. 1998; Omlor et al. 2007; Ball et al. 2008; Muthukumaraswamy 2010). Not only the significant enhancement of power in the high γ-band (>60Hz) (Ball *et al*. 2008; Muthukumaraswamy 2010), as well as strengthened cortico-muscle coherence in the low γ-oscillation (<60Hz) (Brown *et al*. 1998; Omlor *et al*. 2007) were revealed using electrocorticographic and non-invasive EEG recordings, respectively. The γ-oscillation is thought to encode the kinematic parameters for dynamic voluntary finger movement rather than isometric contraction (Muthukumaraswamy 2010). It was further reported in the lower-limb task that the γ-coherence is increased during knee- and ankle-joint movements than the static contraction (Gwin and Ferris 2012). Present results suggest that such a functional role of γ-oscillation can be extended to dynamic control for standing posture. Our findings further indicate the significant differences in iEMG and COPv between postures although there was no significant difference in the γ-connectivity strength. It means that the sway-varying γ-connectivity is a common mechanism across various standings and that the standing balance is not directly associated with its connectivity strength. In the TA muscle for posterior sway, the γ-connectivity was not strengthened in the QS condition, contrary to what was observed in the MG. It may reflect the functional differences between both muscles: the MG activates to counteract the gravitational torque, while the TA is not activated clearly in normal standing.

In addition to the γ-connectivity, the β-connectivity was increased in descending direction at the peaks of COMv. Note that the β-connectivity was strengthened in the QS condition and weakened in the ON condition, particularly in the anterior sway. It means the distinct role between the β and γ-frequency bands and the specificity of the cortical control for the one-leg standing. The task dependency of β-oscillation has been found in the hand and forearm movement in which it only emerged in holding a hand steadily and disappeared with initiating a movement (Kilner et al. 1999; Oya et al. 2020). We suspect this difference is derived from the distinct neural origins. The electrocorticographic study has proposed that γ-oscillation originates in the local cortical source, while β-oscillation is derived from networks that include the inner nucleus, such as the thalamus or basal ganglia (Miller et al. 2007). Thus, we consider that the suppressed β-connectivity in ON and WD conditions reflects the switching of the neural circuit for driving EMG activations. The higher background EMG activities in the ON and WD compared to the QS condition might be supported by the local cortical population rather than reflex activity including the inner nucleus.

In the ascending pathway from the muscle to the sensorimotor region, we discovered that 8−13 Hz was the dominant frequency band. It is widely acknowledged that sensory feedback is critical for the neural mechanism underlying human standing (Peterka 2002). Electrical peripheral nerve stimulation synchronizes the brain and muscle at approximately 10 Hz (Hansen and Nielsen 2004), which corresponds to the rhythm of afferent inputs from muscle spindle discharge (Erimaki and Christakos 2008). Moreover, research suggests that the α-band is a key frequency for intermuscular communication (Kerkman et al. 2020; Laine et al. 2021), facilitating the coordination of afferent feedback for muscle synergy. This evidence shows that the α-band plays a functional role in transmitting sensory information.

Strengthened ascending α-connectivity would improve feedback and give an advantage over standing balance. Our results showed a significantly reduced α-connectivity strength in the ON condition, which may be due to the exertion of larger BGA. Sensory gating during motor execution has been a well-known phenomenon (Chapman et al. 1987; Seki and Fetz 2012). We consider that this mechanism can be driven by muscle activation during standing as the same as limb movement, resulting in the reduction of α-connectivity strength.

### Time-varying corticocortical connectivity

This study examined the time-varying θ-connectivity in 4 different postures and identified the 7 connectivity states, including both posture-relevant and posture-irrelevant components. We concluded that the higher occurrences of the WD and ON conditions imply positive effects on balance recovery (State D), whereas lower occurrences indicate negative relevance (States A and B). Supporting this notion, COPv, a marker of behavioral state, was significantly higher in State D. It is also supported by the distribution of strengthened pairs, which are primarily concentrated in the central region, including the sensorimotor area, of State D and are not found in States A or B. We assumed that the other states, State C, E, and F, are not specific to balance control and are shared beyond the postural differences because the occurrences are nearly identical. Previous research on resting-state brain activity demonstrated linkage of connectivity networks measured by EEG, MEG, and fMRI, both in static and dynamic properties (Deligianni et al. 2014; Allen *et al*. 2018; Wirsich *et al*. 2020). These studies proposed that EEG allows us to capture functional electrophysiological networks in a frequency-dependent manner. The potential of electrophysiological connectivity analysis is followed by a recent EEG study using a virtual reality postural paradigm in which the dynamics of α-connectivity links to sources of the resting-state networks (Aubonnet et al. 2023). Based on these findings, we concluded that multiple networks were simultaneously activated for standing balance.

To validate the contribution of state D to postural recovery, we applied the sway-epoch data to the SVM model based on the time-epoch data. Despite our assumption that the occurrence of State D would increase in sway-poch data, no difference in proportion was found when using time- and sway-epoch data. This suggests that the θ-connectivity around the sensorimotor region is not always facilitated due to posture sway, but may depend on additional factors, such as maximum sway velocity or postural strategies. It may be reflected in individual differences in the predominant state and occurrence. To determine the causal relationship between posture stability and cortical connectivity, future research must challenge a more detailed classification of postural states.

### Static properties of posture-varying corticocortical connectivity

It is widely accepted that cortical involvement is immediately increased due to higher postural demands (Ouchi *et al*. 1999; Solopova *et al*. 2003; Thibault *et al*. 2014; Solis-Escalante *et al*. 2019). The present results advance those implications in terms of electrical physiological connectivity. In the θ-frequency band focused on this study, we found that significant pairs in the static connectivity concentrated again on the midline of the central region. Furthermore, the oscillatory power in the θ-frequency band was modulated in a similar region, consistent with previous research (Hulsdunker *et al*. 2015; Stokkermans *et al*. 2023). The increased θ-connectivity and -power in the frontal and central areas are thought to be induced by errors that need to be controlled under uncertain circumstances (Cavanagh and Frank 2014), and to serve to facilitate sensory-motor processing for standing balance (Mierau *et al*. 2017; Chen et al. 2022). Our findings support these notions and underline the importance of the θ-frequency band for standing balance.

Since standing posture is thought to be achieved through sensory feedback (Peterka 2002), a cortical mechanism for error feedback would be shared with the system in upper limb movement (Scott 2012). The neural mechanism must be supported by a hub function between the local networks, which include the prefrontal, sensorimotor, and occipital regions. In particular, the anterior and midcingulate cortex are likely candidates for these mid-frontal θ-band activities. The cingulate cortex relays cortical and sub-cortical regions and communicates with the frontal and motor cortices to monitor actions and cognitive learning (Brown and Braver 2005; Debener et al. 2005; Womelsdorf et al. 2010; Cavanagh and Frank 2014). In line with those rationales, we assumed that posture-related θ-connectivity originates from the networks including the cingulate cortex, and is projected in the central midline on topography.

For the α- and β-connections, similar significant pairs were found in the topography and connectivity matrix. However, we infer that different information was transmitted across each frequency band. The global α-band network is linked to higher-order cognitive functions, like attention and response selection, which involve the front-parietal network (Hon et al. 2006; Sadaghiani et al. 2012). Research suggests that a reduction in α-synchrony, particularly in the central region, occurs during the preparation phase for voluntary arm movement (Lu et al. 2011) and postural response (Fujio *et al*. 2023), which may serve to maintain a stable internal state for sensory processing. We assume that the tendency of reduced α-connectivity in ON and WD conditions is led by the higher postural demand and contributes to rapid response. On the other hand, β-band cortical connectivity is related to motor execution and learning. It has been proposed that the β-connectivity is primarily found in the motor and parietal cortex and is associated with ongoing movement correction through sensory feedback (Androulidakis et al. 2006; Hillebrand et al. 2012; Chung et al. 2017). The β-connectivity strength increases between interhemispheric sensorimotor areas in a complex bimanual task during the learning phase and decreases afterward (Andres et al. 1999; Gross et al. 2005). In standing posture, we suspect that this connectivity reflects the ongoing postural adjustment which would be required more in the WD condition. Notably, β-connectivity is relatively weak in the ON condition, suggesting the differences in limb coordination between normal and single-leg standing. We interpret the present findings as highlighting the flexibility of corticocortical connectivity in response to both postural demands and movement properties (Lau et al. 2014).

## Conclusion

We investigated sway- and time-varying corticocortical and -muscular connectivity while maintaining postures. The sway-varying corticomuscular connectivity was enhanced at the sway-velocity peak. The predominate frequency was γ- and β-band for the descending connectivity and α-band for the ascending connectivity. For time-varying cortical connectivity, we tracked cortical θ-connectivity dynamics using the time-window method and identified the 7 different connectivity states across 4 different postures. Three of 7 states could be estimated to be posture-relevant states based on the proportion of posture conditions. Finally, the averaged cortical connectivity strength was compared across 4 postural conditions as a static property. The results showed enhanced θ-connectivity in the central region once again, indicating the robustness of this connectivity for standing control. These findings support the previous notion that θ-connectivity promotes sensory processing and standing balance. This study sheds light on the cortical processing of standing posture, particularly its dynamics in response to postural sway and over time.

## Acknowledgment

The authors would like to thank Enago (www.enago.jp) for the English language review.

## Supplementary materials

**Supplementary Figure S1.**
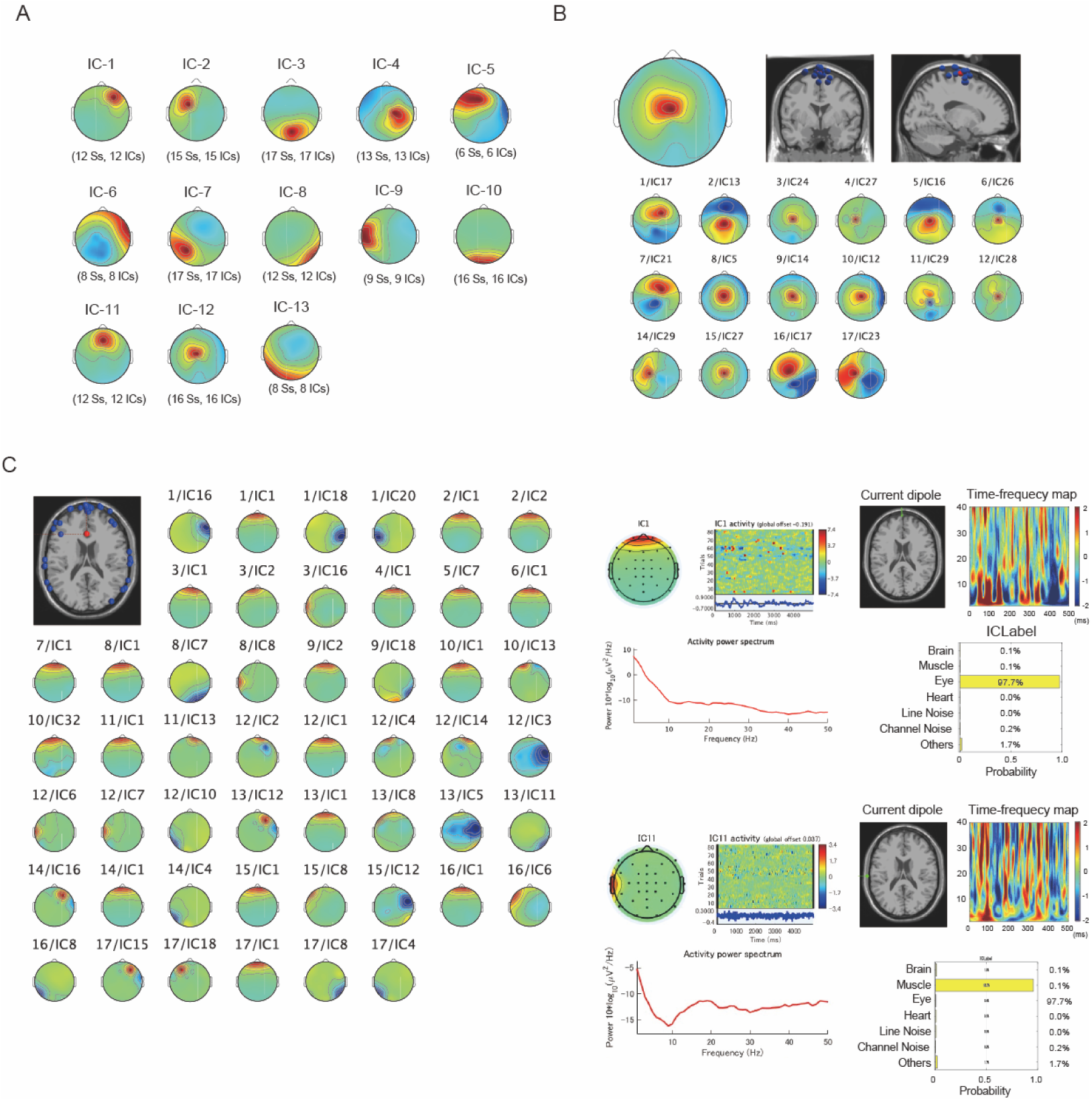
Independent Component Clusters and noise components. A. The 12 clusters contain ICs from more than half of the participants, except for the noise cluster. One IC was chosen from the same subject in each cluster based on the averaged MNI coordination of the equivalent dipole location. B. Thes motor IC was chosen for the corticomuscular connectivity analysis. A single IC was chosen for each participant based on the NMI coordination that was closest to the average position. One participant was not part of the cluster. C. A noise cluster associated with eye blinks and movements was removed from the original EEG time series. We also rejected the ICs for muscle activities using the ICLabel function, which checked the PSD, dipole location, and power distribution in the time-frequency map.

**Supplementary Figure 2.**
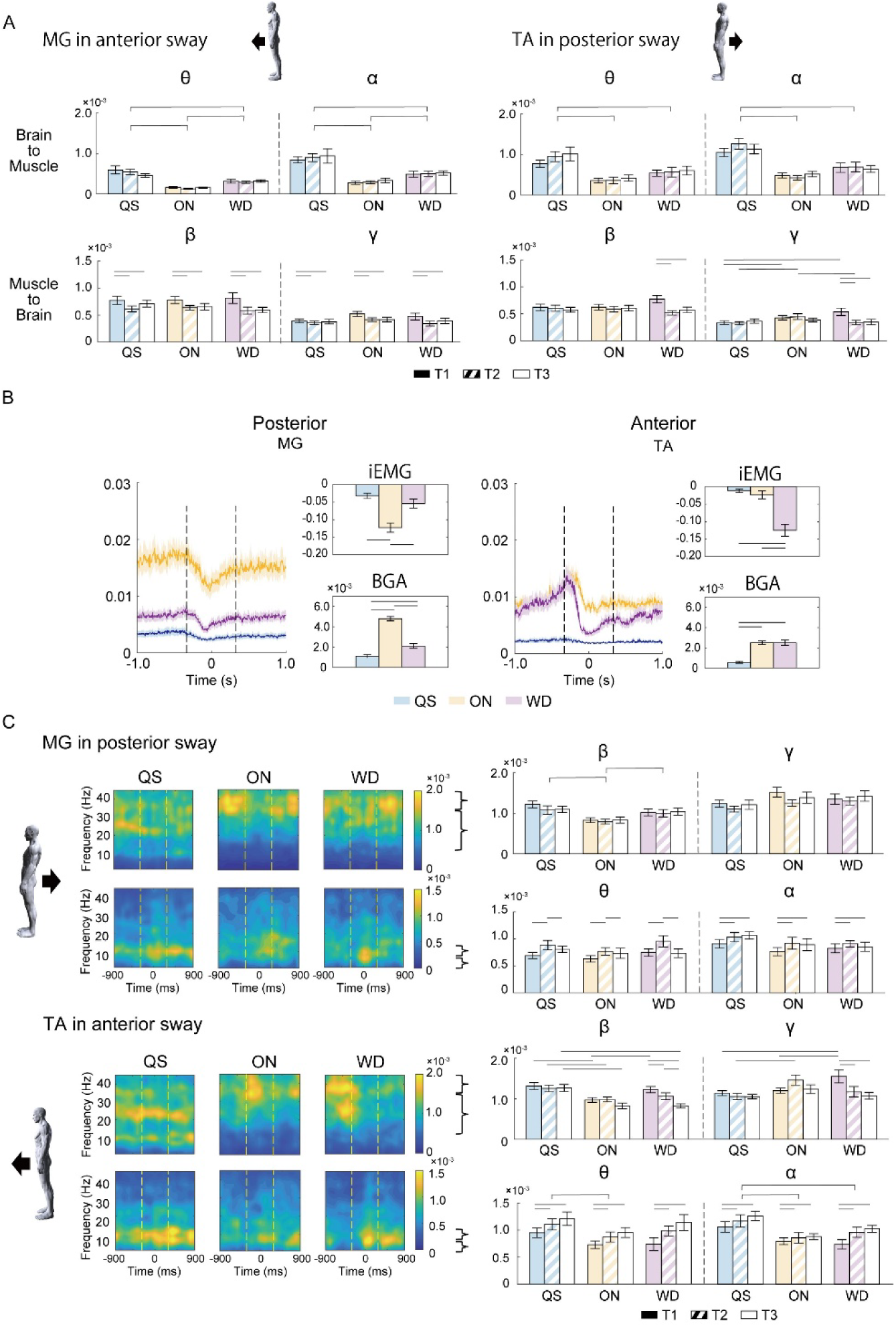
Additional results of the sway-varying cortico-muscular connectivity in the TA and MG. A. The connectivity strength of the corticomuscular connectivity in the θ- and α-bands for the descending direction and that in the β- and γ-bands for the ascending direction. B. EMG activities for 3 standing conditions while swaying anterior (the tibialis anterior muscle; TA) and posterior (MG) directions. C. The sway-varying corticomuscular connectivity in the MG and TA during non-compensatory activity: the MG for posterior sway and the TA for anterior sway. QS; quiet standing, ON; one-leg standing, WD; standing on a piece of wood.

**Supplementary Figure 3.**
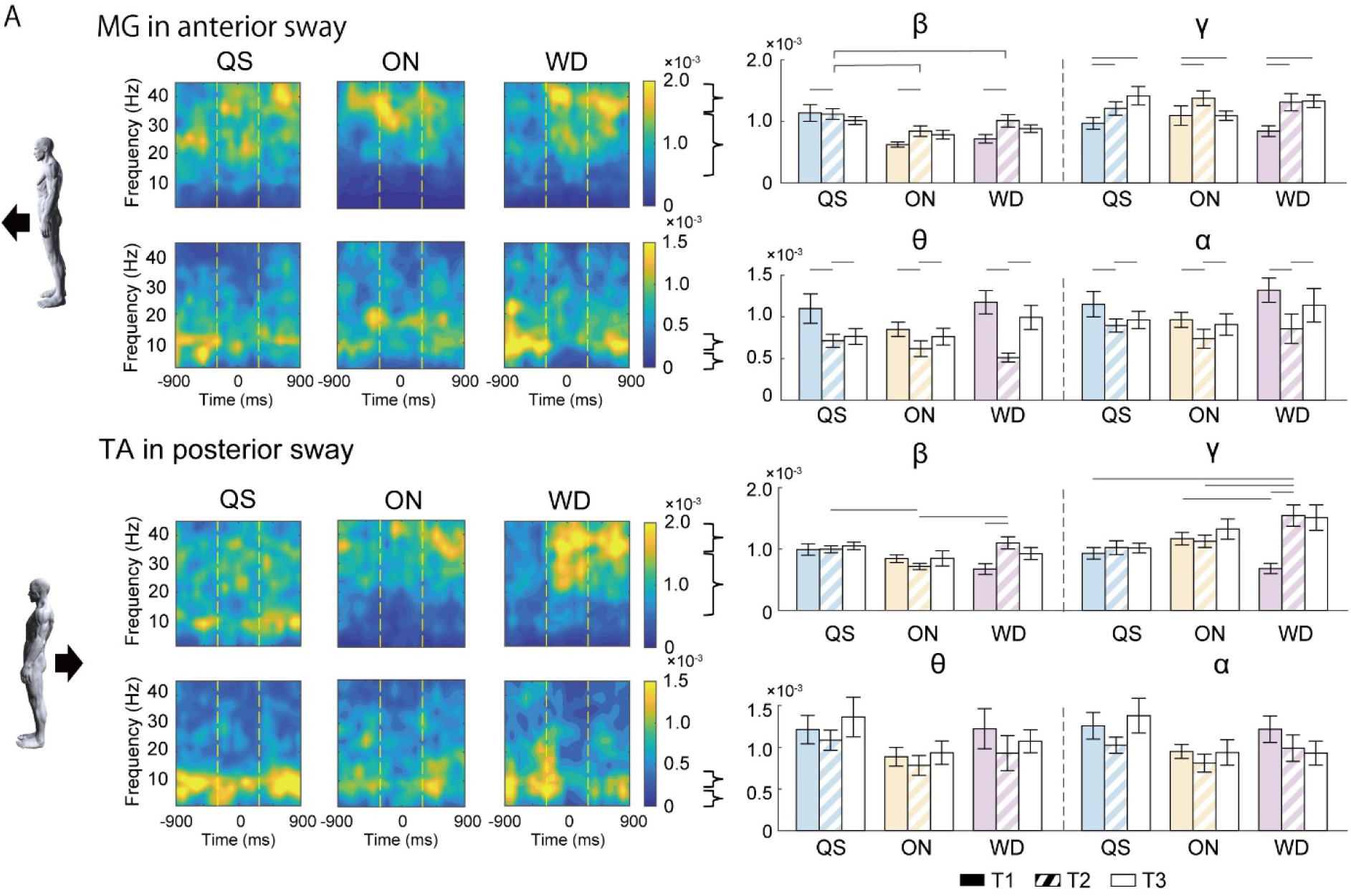
Source level analysis for the sway-varying corticomuscular connectivity and the summary for the other muscles. A. To validate the results in the sensor level analysis, the source level analysis was used for the corticomuscular connectivity. The time-frequency maps and the connectivity strength between the motor-related IC, MG, and TA were shown. In the TA in posterior sway, the main effect was observed in the α-connectivity in the ascending direction, but there is no significant pair in the post-hoc test.

**Supplementary Figure 4.**
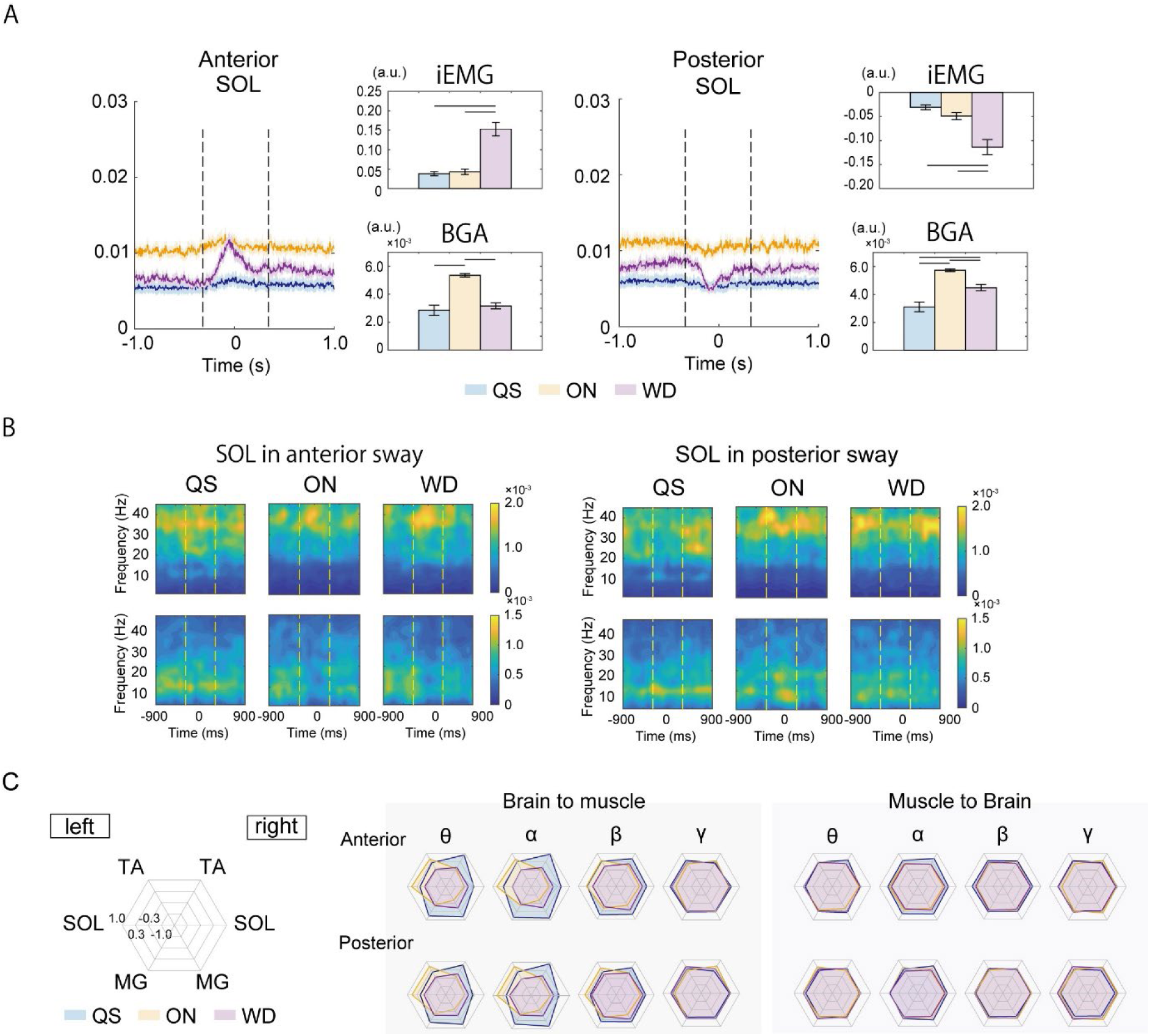
The sway-varying corticomuscular connectivity in the other muscles. A. EMG activities in the right SOL muscle during 3 standing conditions including anterior and posterior sway. B. Time-frequency maps of corticomuscular connectivity in the right SOL muscle. Similar to the MG muscle, the enhancement of the descending communication was observed in β- and γ-frequency bands during EMG activation (T2), particularly in the QS and WD conditions. C. Radar plots shoeing normalized corticomuscular connectivity in both lower-limb muscles. The connectivity strength was normalized under the standing conditions during the T2 period. T1; −900∼300ms, T2; −300∼300ms, T3; 300∼900ms relative to peak COMv. QS; quiet standing, ON; one-leg standing, WD; standing on a piece of wood.

**Supplementary Figure 5.**
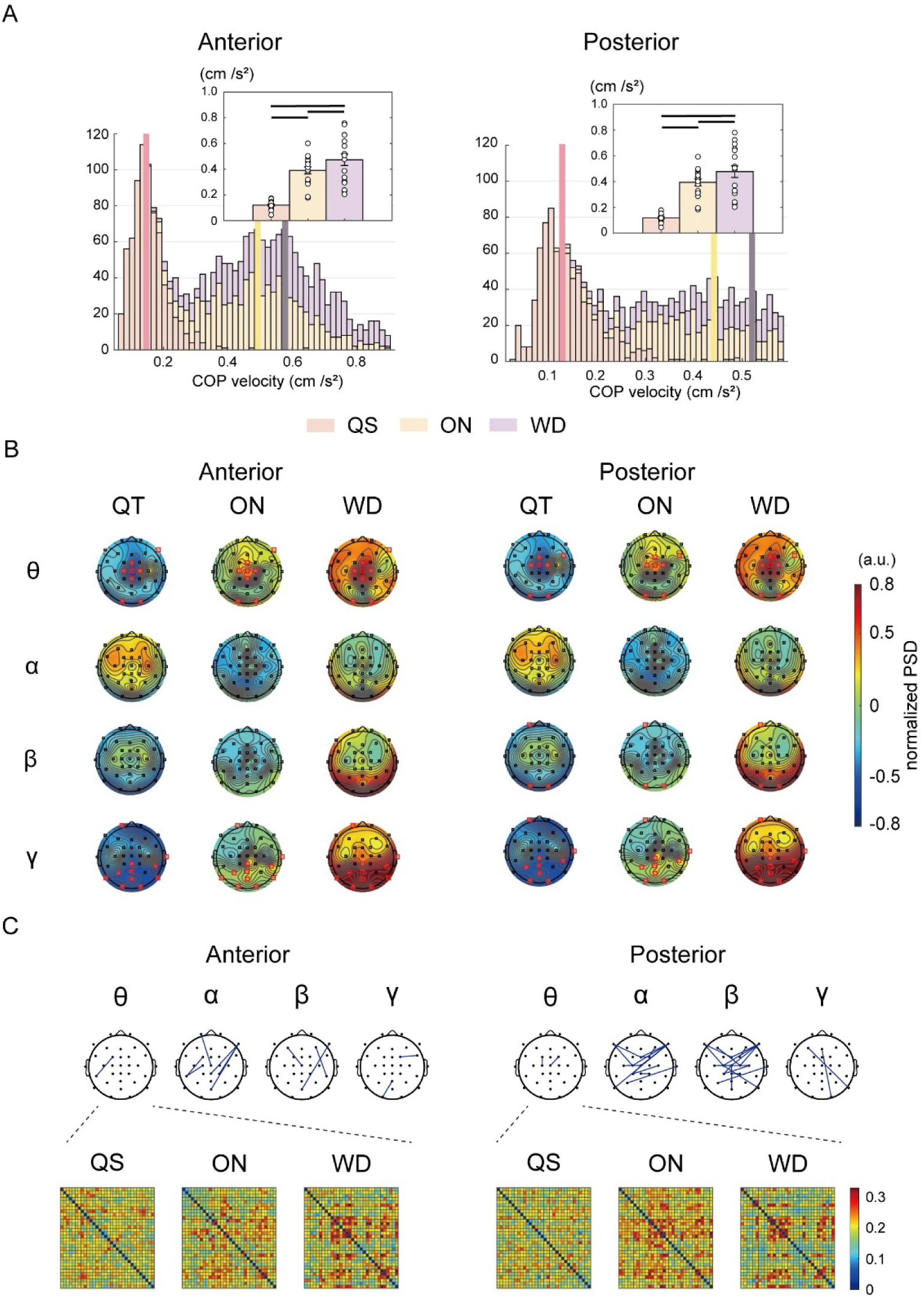
Histogram of COPv, PSD, and corticocortical connectivity in the sway-epoch. A. Histogram of the COPv averaged in swaying anteriorly. Statistical analysis revealed a significantly higher COPv in the WD condition compared to the ON and QS conditions. The COPv was also higher in the ON condition than in the QS condition. B. Histogram of COPv in posterior sway. C. The significant electrodes and distribution of the PSD were depicted on the topography. Similar patterns can be seen in the results of the static properties. PSD in the same time frame for all EEG channels. Statistical tests revealed a similar dispersion of PSD and significant channels on the topography map for both anterior and posterior sway. Significant higher θ-power was observed on the central midline electrodes in ON and WD conditions, while the β- and γ-power was statistically increased primarily on the occipital area. D. Posture-related changes in cortical connectivity were primarily observed at θ-, α-, and β-frequency bands. There were 3.0 ± 2.2 significant pairs in anterior sway, and 8.7 ± 7.9 in posterior sway in each frequency band (anterior, θ: 1, α: 6, β: 3, γ: 2 pairs, posterior, θ: 2, α: 14, β: 17, γ: 2 pairs). The α- and β-connectivity show similar trends with significant changes more noticeable in posterior sway. It implies the directional specificity of the cortical connectivity. ST; sitting on steel, QS; quiet standing, ON; one-leg standing, WD; standing on a piece of wood.

**Supplementary Figure 6.**
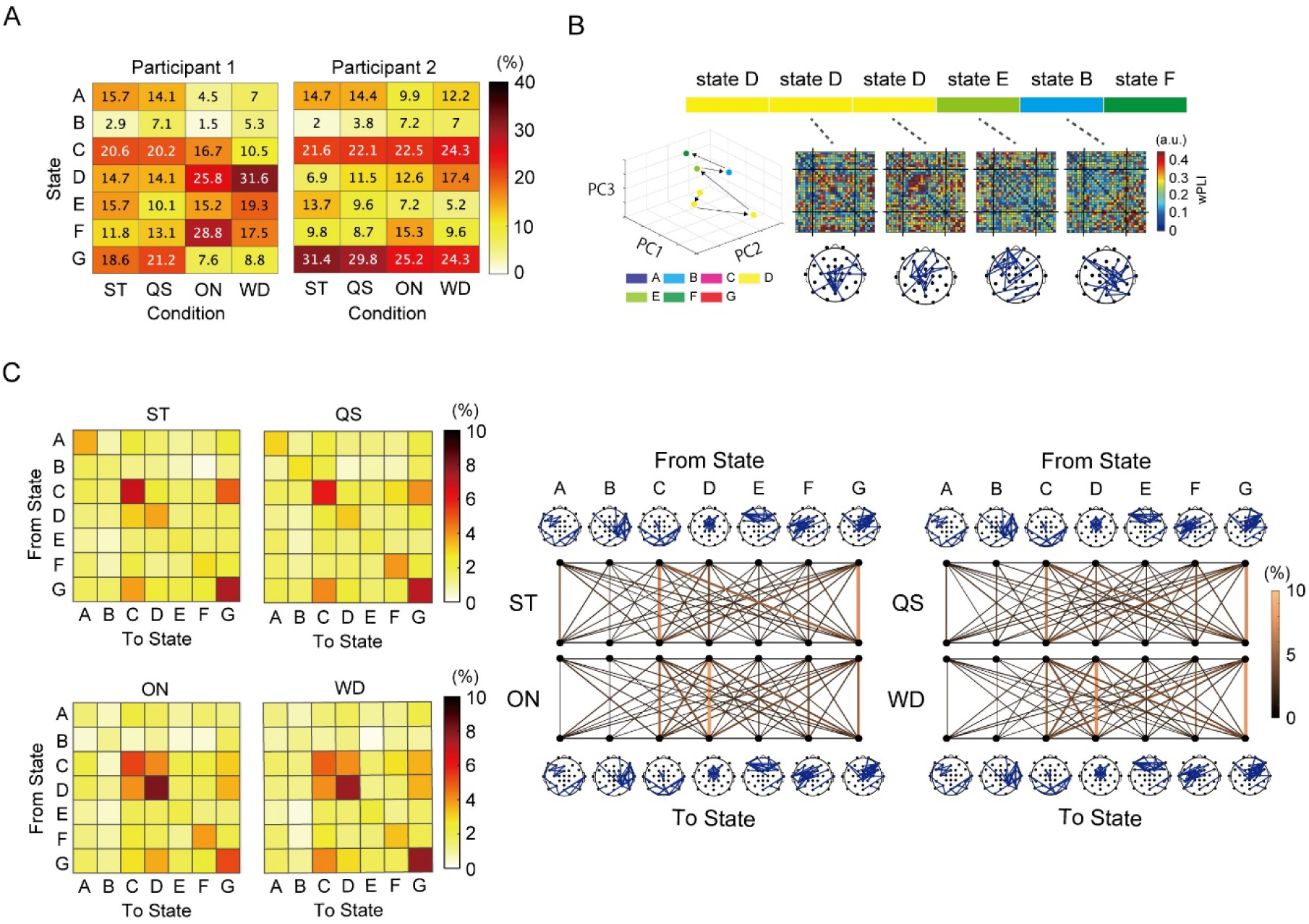
Transitions of the connectivity state over time. A. Examples from two participants for the proportion of postural conditions in 7 connectivity states (State A–G). Regarding the State D, the proportion tends to be equal among QS, ON, and WD in Participant 2, whereas Participant 1 has a typical proportion similar to the group average. B. A shift in connectivity state in a single participant. The feature vector was depicted in 3-dimensional space with arrows indicating the transition direction. The heat maps of the wPLIz for all pairs and the top 5% of strengthened pairs are shown. PC1; principal component, PC2; secondary component, PC3; third component. C. Features of the connectivity transition. The transition matrices in 4 conditions were represented in 2 different ways.

## Figures

Figure 1. Overview of EEG-processing pipeline

Figure 2. Tracking of the EMG activities and cortico-muscular connectivity while postural sway

Figure 3. Time-varying corticocortical connectivity and its differences among postural conditions

Figure 4. Static properties of Cortico-cortical connectivity and PSD

Supplementary Figure S1. Independent Component Clusters and noise component

Supplementary Figure S2. Additional results of the sway-varying cortico-muscular connectivity in the TA and MG.

Supplementary Figure S3. Source level analysis for the sway-varying corticomuscular connectivity and the summary for the other muscles.

Supplementary Figure S4. The sway-varying corticomuscular connectivity in the other muscles.

Supplementary Figure 5. Histogram of COPv, PSD, and corticocortical connectivity in the sway-epoch

